# Two decades of creating drought tolerant maize and underpinning prediction technologies in the US corn-belt: Review and perspectives on the future of crop design

**DOI:** 10.1101/2020.10.29.361337

**Authors:** Carlos D. Messina, Mark Cooper, Graeme L. Hammer, Dan Berning, Ignacio Ciampitti, Randy Clark, Christine Diepenbrock, Carla Gho, Mike Jines, Travis Lee, Ryan McCormick, Eduardo Mihura, Dean Podlich, Jose Rotundo, Matt Smalley, Tom Tang, Sandra Truong, Fred van Eeuwijk

## Abstract

Over the last decade, society witnessed the largest expansion of agricultural land planted with drought tolerant (DT) maize (*Zea mays* L.) Dedicated efforts to drought breeding led to development of DT maize. Here we show that after two decades of sustained breeding efforts the rate of crop improvement under drought is in the range 1.0-1.6% yr^−1^, which is higher than rates (0.7% yr^−1^) reported prior to drought breeding. Prediction technologies that leverage biological understanding and statistical learning to improve upon the quantitative genetics framework will further accelerate genetic gain. A review of published and unpublished analyses conducted on data including 138 breeding populations and 93 environments between 2009 and 2019 demonstrated an average prediction skill (*r*) improvement around 0.2. These methods applied to pre-commercial stages showed accuracies higher that current statistical approaches (0.85 vs. 0.70). Improvement in hybrid and management choice can increase water productivity. Digital gap analyses are applicable at field scale suggesting the possibility of transition from evaluating hybrids to designing genotype x management (GxM) technologies for target cropping systems in drought prone areas. Due to the biocomplexity of drought, research and development efforts should be sustained to advance knowledge and iteratively improve models.

**Highlight:** Crop improvement rate in maize increased after implementation of drought breeding efforts. Harnessing crop, quantitative genetics and gap models will enable the transition from genetic evaluation to crop design.

## Introduction

Over the last decade, society witnessed the largest expansion of agricultural land planted with drought tolerant (DT) maize (*Zea mays* L.) While maize hybrids characterized for superior tolerance to water deficits in the US corn-belt were commercialized over 50 years of breeding, it was recognized that there was an important need to accelerate breeding for DT (Campos et al. 2004, Barker et al. 2005). Dedicated research efforts emerged. Following the commercialization of the AQUAmax* hybrids (AQ herein) in 2011 and the widespread drought event in 2012 (Boyer et al. 2013), the average area of the US corn-belt planted to DT maize hybrids grew quickly to over 20% of the total area (McFadden et al., 2019). In drought prone areas in the western US corn-belt, the land allocated to DT maize can reach 40-60%, as documented for the states of Nebraska and Kansas. Although molecular breeding made feasible the development of most commercial DT products (Cooper et al. 2014a,b) gene editing and transgenic approaches demonstrated the potential for yield improvement under water deficit (Castiglioni et al., 2008; Guo et al., 2014, Habben et al., 2014; Shi et al., 2015; Adee et al., 2016; Shi et al., 2017). Gene edited maize for modified expression of the ARGOS8 gene yielded 33 g m^−2^ more than a control under flowering stress but not grain fill stress (Shi et al., 2017). Similarly, under water deficit maize transformed with ARGOS8 yielded 35 g m^−2^ more than transgene negative hybrids (Shi et al., 2015).

AQUAmax* DT maize is the most studied brand of maize of this class. Over thousands of comparisons and environments in contrasting geographies, AQ maize yielded 37 g m^−2^ more than non-AQ maize when exposed to drought stress. Yield improvement under drought increased with planting density to at least 6.9 pl m^−2^, where the yield difference was 50 g m^−2^ (Gaffney et al., 2015). An important attribute of AQ hybrids is that the yield improvement under water deficit did not come at the expense of reduced performance under irrigation (Hao et al., 2015a,b; Lindsey and Thomison, 2015; Gaffney et al., 2016; Adee et al., 2016; Zhao et al., 2018). This outcome of breeding is consistent with well-defined objectives. The systematic application of 1) selection to a strong legacy germplasm with high levels of drought tolerance (Bruce et al., 2002; Duvick, 2005; Cooper et al., 2014a) evaluated in managed-stress environments and the target population of environments (TPE), 2) precision phenotyping methods, 3) physiological knowledge to inform selections, and 4) advanced predictive analytics, enabled breeders to achieve the objectives (Cooper et al., 2014a,b; Messina et al., 2011). AQ technology was developed for current cropping systems, but increased seeding rates were required for these hybrids to fully express their biological potential (Gaffney et al., 2015; Lindsey and Thomison, 2015). On farm trials followed to demonstrate the advantages of GxM technology (Gaffney et al., 2015). Because of the lower water use of AQ hybrids but maintenance of harvest index (HI) under water deficit (Hao et al., 2015b; Mounce et al., 2016; Zhao et al., 2018), the increased plant population was required to fully utilize the available soil water. Modeling studies using AQ hybrids indicated that reduced stomatal conductance under high VPD (Messina et al., 2015) can increase the water use during the reproductive period at the expense of the vegetative phase. The improved water status during the critical window for kernel set, and the smaller size of the ear at silking (Messina et al., 2011; Messina et al., 2018) can underpin the observed shortened anthesis-silking interval (ASI; Cooper et al., 2014a,b), higher silk number under water deficit (Messina et al., 2019) and the maintenance of harvest index under drought. The experience developing DT maize for the US corn-belt showed how the integration of biological knowledge can increase the rate of crop improvement.

A product development process pipeline in a seed industry is represented in Figure 1. The pipeline starts with the creation of millions of doubled haploids in maize with genotypes that were never tested in the field (Fig.1, 1). Prediction methodologies are utilized to select families and individuals for further testing at all stages during product development. Throughout various stages of testing and selection, the number of individuals tested in field trials reduces to tens of hybrids. Prior to commercialization, these hybrids are evaluated in large areas in thousands of locations (Fig. 1, 2; Gaffney et al., 2015). Around the time of commercialization, agronomists start optimizing the management practices for optimal performance. Further knowledge about the product is gained once the hybrids are grown in farmer fields (Fig. 1, 3). At the early stages of breeding, genotypes are evaluated in few environments, which grow exponentially as these hybrids move through the pipeline. It is not until advanced stages of product evaluation that the norms of reaction and responses to agronomic management are understood. Prediction technologies that account for genotype (G), management (M) and environment (E) were developed to support AQ development to overcome the testing constraints at early stages of development (Cooper et al., 2014a,b; Messina et al., 2018).

**Figure 1.**
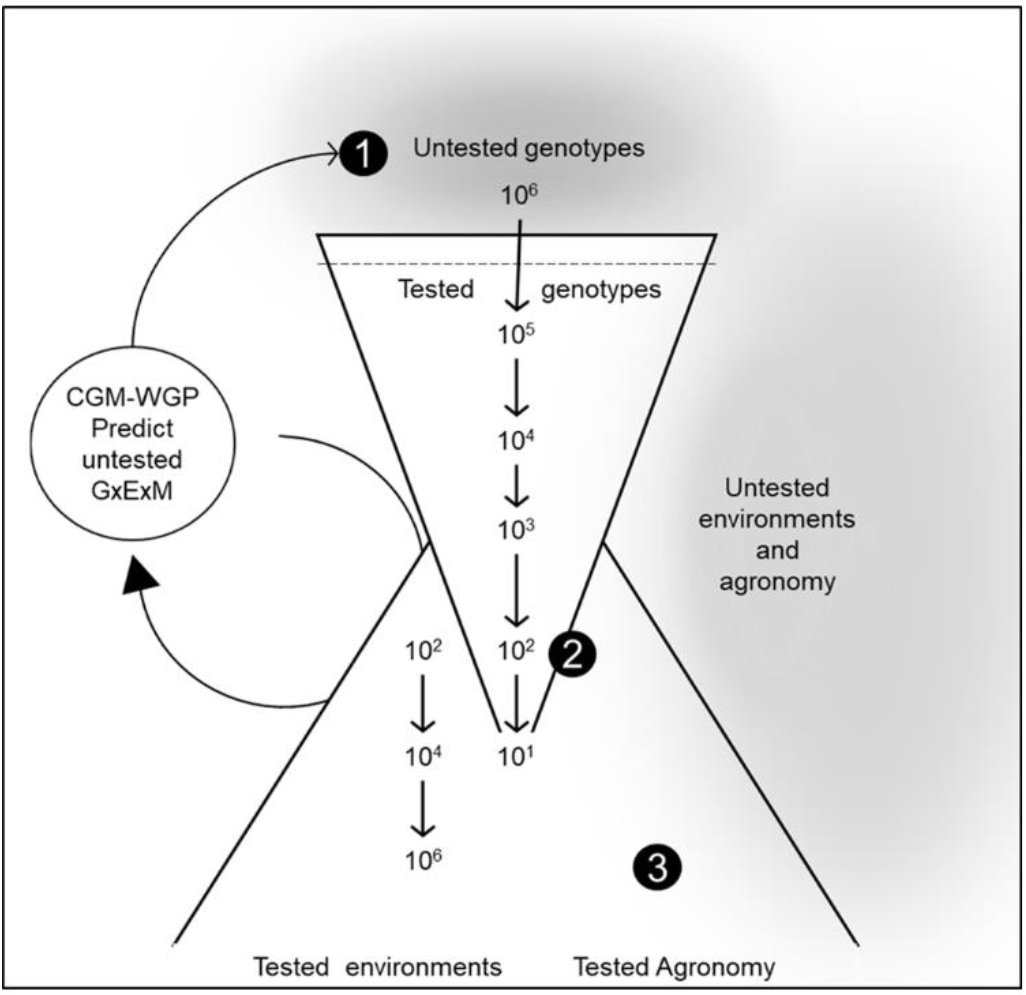
Schematic of a product development process pipeline in a seed industry from the creating of genotypes that were never tested in the field (1) to the time these are testing at scale (2) to the optimization of agronomic practices and growth at farmer fields (3). Crop growth model-whole genome prediction (CGM-WGP) methodology uses statistical learning and biological understanding to prediction and it is built iteratively as more information is gained through the process development pipeline.

Leveraging the experience from developing DT maize hybrids for the US corn-belt, this paper is structured in three parts: Breeding (Fig. 1), Prediction, and Design. Firstly, we review breeding for genetic gain in maize yield under water deficit conditions that has been achieved during the last decade. Secondly, we demonstrate advances in prediction methods, creation and use of DT maize hybrids throughout the US corn-belt. In this section, we discuss prediction applied to the early (Fig.1, 1), precommercial (Fig.1, 2), and on-farm (Fig.1, 3) stages of the hybrid product development pipeline. Thirdly, we put forward a perspective for future application of new design methods that have emerged from the broader complex systems research community in the further development and use of non-transgenic DT maize hybrids. In this section, we discuss leveraging knowledge created at different stages of product development to predict and create GxM technologies at early stages of breeding.

## Breeding

### 1.1 Yield improvement for drought tolerance during two decades of breeding

Over 80 years of reciprocal recurrent selection, breeders increased temperate maize yields at an average rate of 8.6 g m^−2^ yr^−2^ (Cooper et al., 2014b). Because the TPE in the US corn-belt includes various water deficit types (Messina et al., 2015), breeders also improved yields under water deficit at a rate of 6.2 g m^−2^ yr^−1^ (Cooper et al., 2014b). Crop improvement was largely driven by phenotypic selection, with molecular breeding methods contributing to maintain or increase the rate of genetic gain in the first part of the 21^st^ century. After a decade since the introduction of DT maize, it is opportune to ask whether the strategies and technologies put in place in the late 2000s were conducive to increase or maintain the rate of genetic gain.

The comparison between non-AQ and AQ hybrids, and between the first and the new generation of AQ hybrids can provide a first answer to this question. To this end, a set of experiments was conducted in managed-environments in 2019 in Viluco (Chile), Woodland (CA), Garden City (KS), and Plainview (TX); the latter three are in the United States. The first generation AQ includes hybrids P1151, P1498, P0636, P0506 and P0760, all commercialized between 2011 and 2015. The new generation AQ includes P0574, P0657, P0622, P1244, and P1443 commercialized in 2017 and 2018. A set of non-AQ, P0987, P1197, P0801, P1311, P1422, P0789, P1366, P1370, P0950, and P1138, commercialized between 2012 and 2018 were included as a reference of improvements achieved without targeted breeding for DT. The experiment was conducted in four-row plots of 5.2 m of length, and three replicates per location. Irrigation treatments included water withdrawal around flowering and grain filling periods. An irrigated (well-watered) control was included in all locations. Because of the previously reported differential response of AQ (Gaffney et al., 2015; Lindsey and Thomison, 2015; Adee et al., 2016) and other DT maize hybrids (Hao et al., 2019) to plant population, the experiments were grown at 2.5, 4.4, 6.4, 8.4, and 10.4 pl m^−2^. Irrigation quantities for well-watered, flowering stress and grain fill stress were as follows: 854, 741 and 818 mm at Viluco, Chile; 323, 120 and 76 mm at Woodland, CA; 356, 0 and 89 mm at Garden City, KS; and 457, 220 and 276 mm at Plainview, TX.

Results demonstrated that limited-irrigation treatments were effective in reducing yields at optimal seeding across locations. Mean yields across hybrids for flowering stress and grain fill stress were 812 and 937 g m^−2^, respectively. These results contrast with observed yield of 1562 g m^−2^ for a well-watered control (*P*<0.05). Plant population treatments were also effective, with the amplitude of the yield response to population varying between 312 to 625 g m^−2^. The largest differences among hybrid groups were expressed at optimal plant population for yield, which varied by water deficit scenario. The rate of genetic gain of AQ increased with plant population for flowering stress, expressed a definite optimum under grain filling, and showed no clear pattern under well-watered conditions (Fig. 2). At optimal plant population the genetic gain for AQ hybrids was higher (1.0-1.6% yr^−1^) than prior estimates of genetic gain in maize yield under water deficit conditions (0.7% yr^−1^, Cooper et al., 2014a). Under well-watered conditions, the genetic gain for this hybrid set was comparable with prior estimations (Fig. 2; Cooper et al., 2014b).

**Figure 2.**
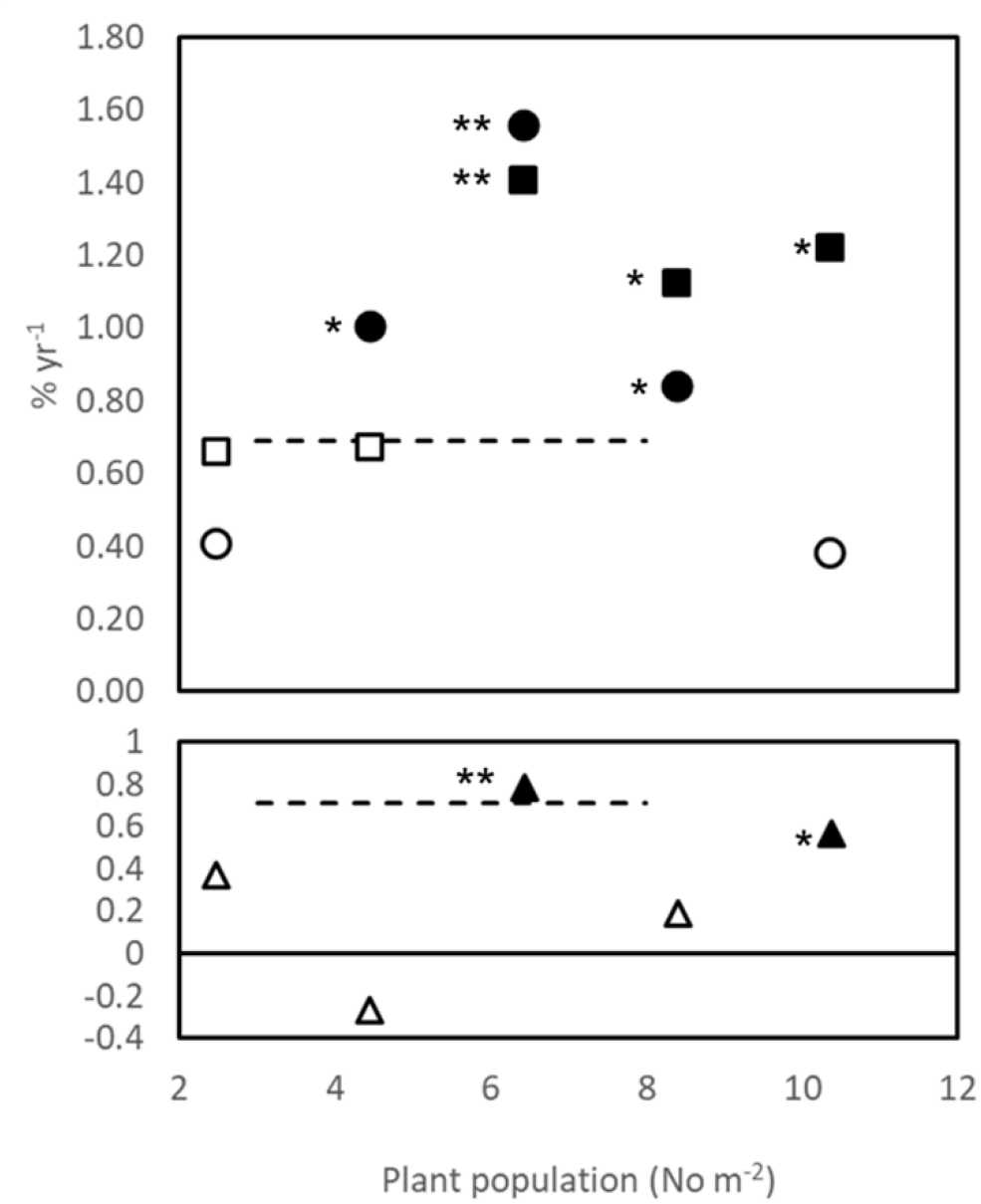
Genetic gain for AQ maize measured in a multienvironment trial across a range of plant populations and with water deficits imposed at flowering time (■) and grain fill (●) stages of development, and a well water control (▲). Genetic gain expressed as the ratio (%) between two cohorts (2011-2015 vs. 2017-2018) of AQ hybrids and average yield for the density and water stress environment combination. Significant differences in yield between cohorts shown as closed symbols (* P<0.1; **P<0.05). Genetic gain for non-AQ hybrids estimated by Cooper et al. (2014a) under water deficit conditions shown as dashed line.

The sustained genetic gain under flowering stress suggests improvement for reproductive resilience. Silk elongation is susceptible to water deficit (Hall et al., 1982; Fuad Hassan et al., 2008; Turc et al., 2016). The reduction in ASI and kernel abortion was demonstrated in prior studies (Edmeades et al., 1993; Bolaños and Edmeades, 1996; Bruce et al., 2002; Messina et al., 2019). Improved kernel set in modern hybrids could be associated with reduced ASI and more silks pollinated, but also to reduced competition within the ear for assimilates among pollinated silks (Messina et al., 2019) and resource availability per kernel (Edmeades et al., 1993; Bolaños and Edmeades, 1996). Under grain filling stress, genetic gain was found to decrease when population increased beyond 6 pl m^−2^ (Fig. 2). This optimum indicates that the observed increased in reproductive resilience under flowering stress treatments extended to grain fill, likely through reduced abortion, but also that limited water availability may have led to an early termination of grain fill limiting the realization of an increased kernel set. Prior studies suggest that yield improvement was not associated with increased water capture at constant density (Reyes et al., 2016; Messina et al., 2020a) and that AQ hybrids rather shifted the patterns of water use instead of increasing total water capture (Cooper et al., 2014a; Messina et al., 2015). A recent study demonstrated that water capture differed between planting density treatments under water deficit conditions but not between double cross and single cross hybrids (Messina et al., 2020a). The absence of a differential genetic gain under well-watered conditions is consistent with the selection criteria focused on improvement of yield under water deficit while not compromising yield potential under well-watered conditions (Gaffney et al., 2015). Taken together, the result supports our current understanding of the mechanisms underpinning genetic gain in maize, which suggest artificial selection did not improve water capture but improved the ability of individual plants to support reproductive structures under crowded stands and stress. Increasing water capture was advocated as a path to improve drought tolerance in maize (Tuberosa et al., 2002; Hammer et al., 2009; Ruta et al., 2010; van Oosterom et al., 2016; Hochholdinger et al., 2018), which seems to remain an unexplored opportunity. These results agree with theoretical predictions using simulation modeling (Cooper et al., 2020b), thus creating the opportunity to use prediction methodologies to hasten crop improvement (Cooper et al., 2020a,c).

### 1.2 Strategies for yield improvement under drought

The objective of commercial breeding programs is to make the highest possible rate of genetic gain for one or more traits at the minimum cost (Cooper et al., 2014b, Ramirez-Villegas et al., 2020). Breeding objectives for maize in the US corn-belt generally include yield improvement, drought tolerance, standability including ear and plant height, disease tolerance, and incorporation of transgenic traits for insect and herbicide resistance, among others. A general structure of a breeding program and the hybrid development pipeline is described in Cooper et al. (2014b). Briefly, in hybrid crops such as maize, a drought breeding program is the result of running two breeding programs in parallel that complement each other. Traits conferring drought tolerance may be contributed from lines identified in the female or male heterotic group, or as the result of heterosis (Barker et al. 2005, van Eeuwijk et al. 2010). At industrial scale, a drought breeding program is more complex resulting from the integration of multiple breeding programs. In this case, adaptive traits could be contributed from any of the active programs for temperate maize (Cooper et al., 2014b). Once DT maize lines are identified, further testing occurs to identify superior hybrid combinations. During the testing and advancement of hybrids, one of more transgenes are introgressed into the parental lines of the hybrids prior to their commercial release. The complexity of industrial programs opens opportunities to optimize processes that minimize costs and increase rates of genetic gain.

Design of breeding strategies and the product development pipeline from the creation of genotypes to the optimization of agronomic practices are promising areas to increase the efficiency and effectiveness of drought breeding. Selection criteria and intensity, the structure of reference populations, and the testing systems implemented to express standing genetic variation for adaptive traits are all decisions breeders make to systematically change the frequencies of the alleles of the genes underpinning adaptation to drought (Cooper et al., 2020a). Modeling and simulation can help breeders manage this complex system and increase the probability of shifting allele frequencies towards the desired directions. However, simulation of biological systems using principles of quantitative genetics is not a new concept and dates back to the 1970s (Fraser and Burnell 1970). The development of simulation software (e.g., QU-GENE software Podlich and Cooper 1998) made more accessible the application of simulation to the study and optimization of breeding strategies. The link functions connecting genotype and phenotype was based on the infinitesimal quantitative genetic models (Lynch and Walsh, 1998; Walsh and Lynch 2018; Cooper et al., 2020a). Using the flexible model *E*(*NK*) to simulate GxE and GxG interactions, Cooper et al. (2005) demonstrated that the largest impact of molecular breeding would be for complex traits such as drought tolerance, where such interactions are commonplace. Extension of this study was demonstrated for breeding for drought tolerance in both maize and sorghum (Chapman et al., 2003; Messina et al., 2011). Studies of maize breeding outcomes revealed behaviors consistent with those of complex systems, including sensitivity to initial conditions, such as the selected set of founder genotypes, and the physiological state of the breeding germplasm (Messina et al., 2011). The accessibility of trait combination associated with peaks of high yield under drought stress were dependent on the distribution of reproductive resilience and canopy architecture in the reference population of genotypes of the breeding program. These studies were possible by nesting the *E*(*NK*) model within a new gene-to-phenotype link function, now represented by hierarchical structure of crop growth models (Messina et al., 2011). These studies both indicated that the largest opportunities for improving genetic gain under drought stress were whole-genome prediction enhanced by design of field testing and evaluation strategies based on biological insights. Uncertainty determined by the linkage disequilibrium conditioned by founder genotypes, the stochasticity of recombination, environment, and internal variation of the system was embraced and incorporated in the decision making. Outcomes from simulation (Messina et al., 2011) implementing concepts for weighted selection (Podlich et al., 1999) informed selection decisions on sampling of the TPE, irrigation protocols for managed-stress environments, and precision phenotyping (Cooper et al., 2014a; Cooper et al., 2014b). Biological insights were used to design experimental management strategies in key environments to expose genetic variation for adaptive traits. For example, managed-stress environments were implemented in Woodland, CA and Viluco, Chile to expose the germplasm to conditions that were conducive to express variation for traits that affect the dynamics of the water balance, water capture, and reproductive resilience. In shallow soils, traits such as limited transpiration (Choudhary et al., 2013; Shekoofa et al., 2015; Tardieu et al., 2017), canopy expansion (Lacube et al., 2017) and silk elongation response to water deficit (Cooper et al., 2014a; Fuad-Hassan, 2018; Messina et al., 2019) would confer adaptation to drought. In contrast, in deep soils, breeding lines with deep root systems can manifest higher yields (Reyes et al., 2015; Messina et al., 2020a). A robust breeding strategy must include selection in both of these environment types to enable identification of trait combinations that contribute to stability of yield performance across the diverse range of environments expected in the TPE of the US corn-belt (e.g., Gaffney et al. 2015). The application of the robust quantitative genetic framework (Lynch and Walsh, 2018, Walsh and Lynch 2018), biological knowledge and precision phenotyping led to the observed large impact on genetic gain (Fig.2).

## Prediction

### 2.1 Progress and application of prediction at early stages of breeding for drought

The application of gene-to-phenotype prediction methodologies, mainly whole genome prediction (WGP), enabled the revolution in molecular breeding (Meuwissen et al., 2001; Gianola et al., 2009; Cooper et al., 2014b; Heslot et al., 2014; Crossa et al., 2017; Voss-Fels et al. 2019). This transformation in breeding was only possible because of the convergence of molecular approaches with other technologies such as double haploid production, and precision phenotyping (Cooper et al., 2014b). These technologies are applied routinely at early stages of breeding programs to enable the generation of and selection upon large numbers of untested and tested individuals increasing the size of the breeding programs (Fig. 1; Araus et al., 2018; Hammer et al., 2019; Washburn et al., 2020). However, ubiquitous GxE interactions under water-limited conditions place a cap on the rate of attainable genetic gain (Voss-Fels et al., 2019; Cooper et al., 2020a).

Transdisciplinary approaches that leverage biological insights and statistical learning methods are changing the ways in which we approach crop improvement (Hammer et al., 2019; Messina et al., 2020b). The challenge to prediction that stems from the need to predict GxExM interactions motivated modeling GxE within statistical frameworks (Jarquin et al., 2014; Jarquin et al., 2017; Li et al., 2018; Millet et al., 2019). Although these statistical approaches are essentially static in character, they can capture the dynamics of crop systems when biological understanding is leveraged in the selection of environmental covariates and the aggregation of information by stages of development known to be of critical importance for yield determination (Millet et al., 2019; Bustos-Korts et al., 2019a,b). Other approaches fully incorporate the dynamics of the crop system. The integration of WGP with crop growth models (CGM-WGP, Technow et al., 2015; Cooper et al., 2016; Messina et al., 2018) is such an example. This approach enables a new generation of prediction methods that, by explicitly modeling GxMxE interactions, has the potential to increase predictive skill and expands domains of inference in both the environmental and agronomic management dimensions. CGM can predict phenotypes for a given genotype and management for productivity and water use, nitrogen loss, and other metrics that can enable decision makers to assess the value of genotypes in the context of environment sustainability (Peng et al., 2020) and contribution to the implementation of a circular economy. Because physiological traits in CGM-WGP are directly modeled using marker information, it is possible to estimate these with accuracies that are dependent on the degree of relatedness between populations to generate prior knowledge, and the genotypes of interest. Physiological traits are parameters in the model that quantify, for example, how transpiration is converted to mass (Tanner and Sinclair, 1983). The stringency of experimental designs and information management increases but the field experimentation demands decrease. In CGM-WGP it is not necessary to measure any physiological traits. However, it is critical to expose the germplasm to environments that elicit trait x environment interactions to enable the estimation of parameters (Messina et al., 2018). When possible, observation of trait physiology is preferred to complement and evaluate estimation approaches. Whether some traits are measured or estimated, CGM-WGP enables breeders to access biological knowledge, physiological and genetic, to inform selection decisions at early stages of breeding when phenotyping of physiological traits is limited at an industrial scale. Advances in high throughput phenomics (Araus and Cairns, 2014; Araus et al., 2018; Reynolds et al., 2020), our understanding of how trait and state phenotypes are connected within modeling frameworks (Cooper et al., 2014b; van Eeuwijk et al., 2019), and the possibility to assimilate phenomics and genomics information within CGM-WGP (Messina et al., 2018) will increase our understanding of adaptation to drought and predictability thereof.

The central hypothesis underlying CGM-WGP is that by harnessing biophysical knowledge through the CGM to capture the gene-to-phenotype relationships for traits contributing to yield variation and consequently trait-by-environment norms of reaction, it is possible to a) understand effects of allele substitution and genetic variation for traits across environments, and b) increase predictive skill. The first demonstration of a reduction to practice of the method (Cooper et al., 2016) used a CGM to enable WGP to predict GxE for yield for one population and two drought environments. Subsequent implementations used Bayes A as a baseline model, which does not model GxE interactions (Messina et al., 2018). The augmented model, CGM-WGP, uses a hierarchical Bayesian algorithm to model the relationship between markers and physiological traits, and the relationship between environment and yield conditioned on agronomic management through the CGM. The comparison between Bayes A and CGM-WGP is an estimator for the capacity of CGM-WGP to model GxExM interactions. CGM-WGP improvement of predictive skill relative to WGP can depend on the similarities between environments, how the environment elicits genetic variation in adaptive traits, and the physiological mechanisms underpinning adaptation to drought.

Results from Cooper et al. (2016) demonstrated empirical application of CGM-WGP for a drought study where there was little improvement of over genomic BLUP alone (Fig. 3). The drought environments considered by Cooper et al. (2016) discriminated the germplasm in a very similar manner. There was a high genetic correlation (*r*_G_=0.88) for yield between the two flowering stress environments included in their study. While the timing of water deficit varied between the two treatments, the same physiological mechanism underpinned the observed tolerance to drought, limiting the expression of differential GxE for yield (Cooper et al., 2016). In contrast, significant improvements in predictive skill of CGM-WGP over WGP alone were observed when contrasting environments (deficit irrigation and full irrigation) and populations expressing contrasting genetic correlations (−.08-0.49) were considered (Fig. 3, Messina et al., 2018).

**Figure 3.**
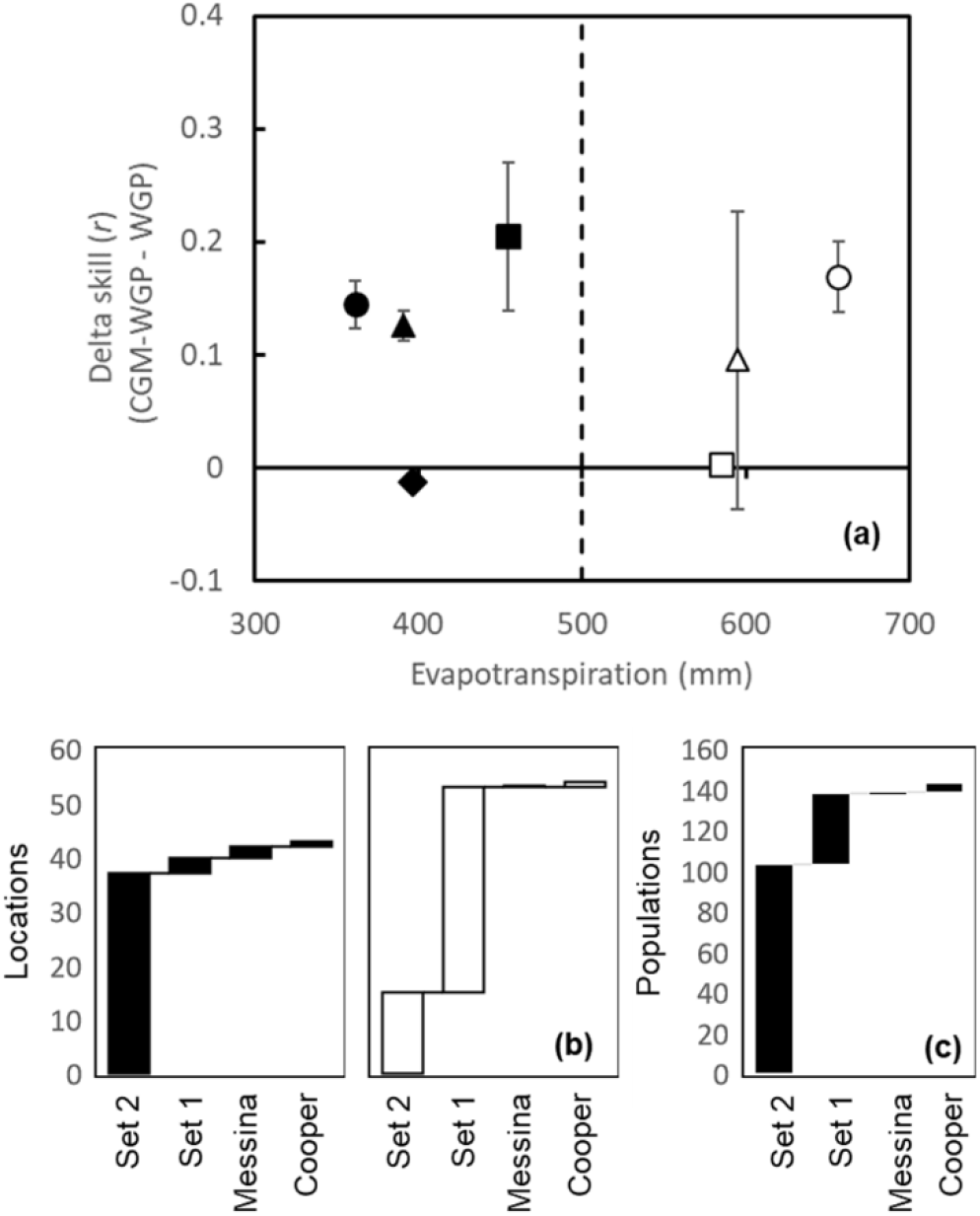
Average prediction accuracy difference between the Crop Growth Model – Whole Genome Prediction methodology and Bayes A as a function of evapotranspiration (A), and number of locations (B) and populations (C) included in each study. Accuracy estimated by the correlation coefficient (*r*) for the validation set. Open symbols indicate results from experiments conducted under irrigation or well-watered conditions in US corn belt. Closed symbols indicated results under water deficit. Mean and standard error of the mean for prior studies: Cooper et al. (2016) (◆), Messina et al. (2018) (▲), Set1 (■), and set 2 (●).

Because adaptation to drought is complex and prior studies seeking to understand and assess the CGM-WGP methodology include few environments and populations (Fig. 3), the potential to improve predictive skill was not fully explored. In the current paper we use larger datasets to further understand the domains of application of CGM-WGP. The hierarchical Bayesian algorithm is from Messina et al. (2018). The maize model (Messina et al., 2015; Cooper et al., 2016) was modified to simulate cohorts of silks, the dependence of silk number on the number of rings per ear and kernels per ring. Silk elongation was simulated as a function of water deficit, and time to silking was determined by both the rate of elongation and the average distance between the cob and the tip of the husk (Messina et al., 2019). The simulated total number of silks determined the attainable harvest index as described in Cooper et al. (2016). The selection of traits to model as a function of markers was informed by assessing the relative contribution to predictability 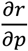, where *r* is the correlation coefficient and estimator of predictive skill and *p* is the variation in the physiological trait (Table 1).

**Table 1.** Sets and values used to define prior distributions for physiological traits that define parameters in the crop growth model.

| Data set | Set 1 |  | Set 2 |  |
| --- | --- | --- | --- | --- |
| | $\bar{x}$ | $\sigma$ | $\bar{x}$ | $\sigma$ |
| Number of rings per ear | 45.0 | 2.6 | 45.0 | 2.6 |
| Leaf appearance rate (°C) |  |  | 0.00275 | 0.00012 |
| Husk Length (mm) | 190.0 | 10.2 | 190.0 | 10.2 |
| Rooting rate (mm day<sup>-1</sup>) | 25.0 | 2.6 |  |  |
| Senescence response to water supply/demand | 0.05 | 0.01 |  |  |

Two datasets were utilized in this study. The first set (S1) included experiments conducted in 17, 12 and 6 locations in the US corn belt in 2017, 2018, and 2019, respectively. A second set of experiments was conducted in Corteva managed-stress environments located in Viluco, Chile and Woodland, CA in 2017 and 2018, where irrigation was managed to satisfy all water demands, and to impose water deficits around flowering, and during grain filling. Irrigation was applied using buried drip tape (Cooper et al., 2016). The genetic material included in both experiments was from crosses between nine non-stiff stalk inbred parents in a half-diallel mating design, and the doubled haploids were crossed to a common tester. A total of 35 families were included in the study. The second dataset (S2) included experiments conducted in managed-stress environments (five to eight locations/treatments per year) with irrigation treatments described above, and in the US corn-belt (one to four locations per year in the states of Iowa and Nebraska). The experiment was conducted between 2009 and 2014 in two-row plots 5.25 m of length for a total of 52 unique location/environment combinations. Hybrids evaluated in the MET shared a common tester within year but not across years. Populations differed among years for a total of 103 populations. Experiments conducted under water deficit included two replications. A set of common checks were included across experiments that enabled a combined analysis across years. Phenotypic analyses of both data sets were conducted applying a mixed-model framework with spatial adjustment for row and columns. Best linear unbiased estimators (BLUEs) by location were used to train both WGP and CGM-WGP for S1, and best linear unbiased predictions (BLUPs) for S2. Prediction algorithms were trained using data from all environments and a sample of 250 genotypes. Predictions were contrasted with observations from genotypes not included in the training set, which comprised 90% of the individuals for S1, and 78% on average per year for S2 (note populations varied with year in S2).

Results from the analyses of S1 and S2 data showed that CGM-WGP is an effective method to model GxE. The experiments encompass environments that range from 250 to 800 mm of evapotranspiration (ET), which covers most of the yield range simulated and observed by Cooper et al. (2020a). It is apparent that the improvement in prediction accuracy is larger under drought conditions, where CGM-WGP consistently improved predictive skill over Bayes A (Fig. 3). More variable results were observed when ET was greater than 500 mm. The estimates of predictive ability for the reference method, Bayes A in this case, were 0.40 and 0.14 for S1 and S2, respectively. The low predictability in S2 was in part due to the presence of significant GxE interactions. Cooper et al. (2016) and Messina et al. (2018) showed genetic correlations for a subset of populations ranging from −0.18 to 0.88. The improvement in predictive skill between CGM-WGP with respect to the base ranges between 0.1 and 0.2 across 103 breeding populations in S2 (Fig. 3). The use of the biophysical model to model GxE improved the estimation of allele values and value of genotype. Considering the following form of the breeder’s equation *R* = *ir_a_σ_a_*, where *R* is response to selection, *σ_a_* additive genetic variance and *i* intensity of selection (Lynch and Walsh, 1998; Voss-Fels et al., 2019), doubling predictive ability (*r_a_*) implies doubling the response to selection or gain under drought stress conditions.

Cooper et al. (2020a) shows that the ratio between the genetic variance 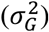 relative to the GxExM variance 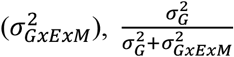, decreases with increasing water deficit. Based on this theoretical work, it is possible to predict that the difference between CGM-WGP would be higher under water deficit conditions than under well-watered conditions. Based on the analyses of 138 populations and 97 environments (Messina et al., 2018; Fig. 3) it is possible to conclude that the combination of biological understanding and statistical learning methodologies can improve predictive skill and therefore will hasten the rate of genetic gain, at least for maize in the US corn-belt.

The sensitivity analyses conducted on the physiological traits for their marginal contribution to predictive skill proved useful to increase prediction accuracy. The uses of optimization for estimation of parameters in biological models is an active and promising area of research (Pathak et al., 2007; Casadebaig et al., 2016; Wallach et al., 2019). Methods for automated selection of traits for a given set of experiments can become an enabler such that breeders can utilize physiological understanding to inform breeding decisions without requiring detailed knowledge of the inner workings of the physiological model.

### 2.2 Delineating areas of adaptation using ex-ante assessment of genotype x management interactions

A critical component of the hybrid maize development process, especially for the improvement of drought tolerance, is the wide area testing of hybrids at farm scale (Fig. 1, 2) and the need to conduct ex-ante analyses of the performance of candidate hybrids for production in the TPE (Cooper et al., 2014; Kruseman et al., 2020). This is necessary because of the need to characterize and manage in the best possible ways ubiquitous GxExM interactions that drive performance in the US corn-belt (Cooper et al. 2014b; Cooper et al., 2020b) and the probability of making incorrect selections and pairings of G and M and the corresponding poor performance and farmer’s risk increase with decreasing 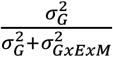. On-farm testing of G and M pairings is the current solution to deal with this problem, but it is expensive and poses a great challenge to breeders and agronomists because of the combinatorial nature of the problem defined by GxExM interactions.

Predicting G norms of reaction for GxExM has been a long-term ambition of both breeders and agronomists. Such tools can enable the placement of hybrids in the geographies, landscapes and fields with customized management to the genetics for famers to realize the biological and environmental potentials of their on-farm systems. But the availability of such tools at scale has been elusive. While tools such as crop models that enable management of irrigation have been around for more than half a century (Jones et al., 2017), their adoption has been low. Only less than 1% of US farmers use these prediction methods to manage irrigation in the United States (United States Department of Agriculture, 2019). While the lack of access to the technology in simplified user-friendly forms can explain the low adoption, particularly in developing countries (Lowenberg-DeBoer and Erickson, 2019), the lack of knowledge on the physiological basis of adaptation and access to genotype-specific information and prescriptions is another probable cause. While efforts have been proposed to use molecular markers to characterize genotypes to enable biophysical prediction (Messina et al., 2006; Yin et al., 2004; Bogard et al., 2020), these approaches have not been applied at scale to predict complex phenotypes.

During the last decade, a method comprised of field experimentation and use of CGM-WGP was developed. Between 2017 and 2019, field experiments were conducted at five to seven locations per year in the US corn-belt, and Corteva managed-stress environments in Woodland, CA and Viluco, Chile. The experiments included variable irrigation interrupted at flowering and grain fill stages or reduced by 25, 50 and 75% of reference ET during the growing cycle. Nitrogen fertilizer rates ranged between 0 and 210 kg ha^−1^ and plant population was reduced and increased by 20 % relative to normal seeding rates for the location. A set of traits were measured based on knowledge of genetic variation of the germplasm as it was developed through the breeding pipeline (Cooper et al., 2014b; Messina et al., 2018). This set included leaf number counts at regular intervals ranging between 3 and 10 days, size of the largest leaf within the canopy (Cooper et al., 2016), ear size at silking (Cooper et al., 2014b), flowering notes, yield and yield components. Carbon assimilation response to light intensity, kernel growth rates, mass during the growth cycle, light interception, specific leaf N of the largest leaf (DeBruin et al., 2013), and transpiration response to VPD (Choudhary et al., 2013; Shekoofa et al., 2015) were measured in a subsample of hybrids to characterize prior distributions (Messina et al., 2018). Parameters within the crop model such as radiation use efficiency and its maintenance at low water potentials, root elongation rates (Reyes et al., 2015; van Oosterom et al., 2016) and other traits for which information was generated to estimate prior distributions, were then estimated using marker information and the procedure described by Messina et al. (2018).

The result of combining a field research program with the use of markers and a prediction methodology to estimate parameters within the crop model was a scalable capability to predict hybrid performance. The data generated through simulation are complementary and thus augment the data collected within the multienvironment trials (MET; Fig. 4a). To test the predictability of the data collected within the MET, these data were segmented through a series of concentric circles with origin placed at a research station in Windfall, Indiana. A subset of the data within the circle was retained as the truth, and the reminder was used to estimate the correlation between the two samples. A second correlation was estimated by averaging the sample of observed values and the simulated values for the location at the origin. The procedure was repeated 50 times to estimate variability of the prediction. For the example shown in Figure 4, augmenting the observed data with predictions based on biophysical knowledge can increase predictive skill up to 300 km when the observed data is at least 50% of the original data. However, this distance could be influenced by the genetic correlation between the central location and the rest of the locations in the region. The improvement in predictive skill of the combined approach (statistical and CGM prediction) increased with predictive skill of the CGM (Fig. 4b). Both methods are capturing different genetic signal that could be utilized to further increase yield prediction through ensemble methodologies (Wallach et al., 2019; McCormick et al., 2020). This result indicates that the approach is predictive, scalable and can both increase predictive skill of statistical methodology while reducing the experimental footprint.

**Figure 4.**
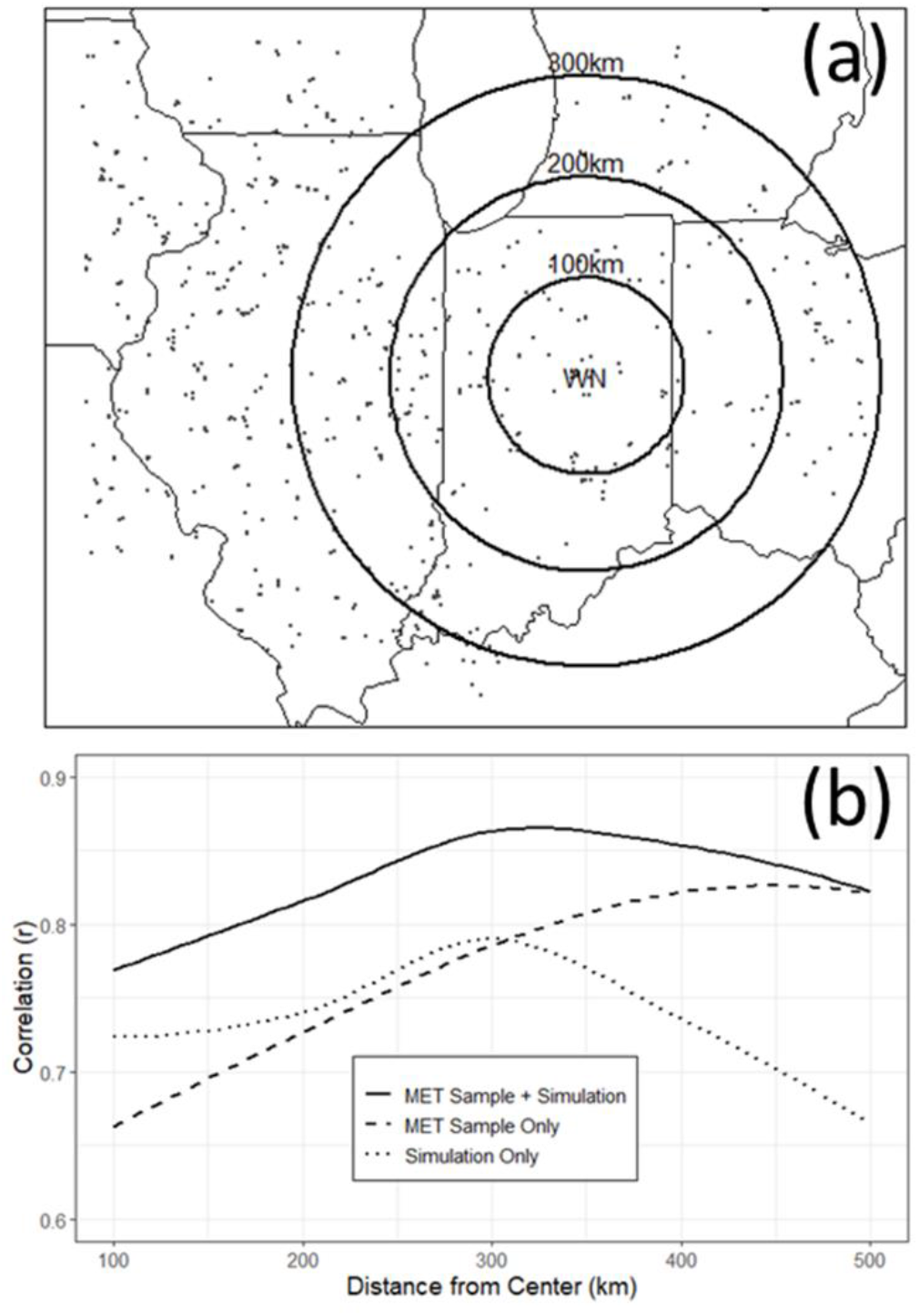
Multienvironment trial for maize (a) and predictive skill variation with distance to the center WN (b). Circles drawn to illustrate areas with constant distant to center WN that define a set of data to calculate predictive skill. Predictive skill calculated for three cases: 1) estimation of accuracy through resampling the data in the multienvironment trial (MET sample only), 2) accuracy estimated by the correlation between crop model simulation and observation (simulation only), and 3) correlation between the average prediction of 1 and 2, with an independent sample of yield data for all the hybrids included in the trial. The correlation (*r*) is calculated across genotypes.

Because the crop models used in this approach to prediction encapsulate biological knowledge (Jones et al., 2017; Hammer et al., 2019), it is possible to utilize biological insight to understand the physiological basis of adaptation and yield determination under various environments. Figure 5 shows the relative contribution of different traits to yield determination within a gradient of yield. To determine the sensitivity of yield to variation in physiological traits, the crop model was run for 30.5 million simulations (137 hybrids over 54 historical years across production regions of the US) comprised by changes in management and environment. For this study, weather and soil data were from National Oceanographic Administration and Natural Resources Conservation Service, respectively (see details in Messina et al., 2015). The graphical representation of the contribution of various physiological traits to yield shows that biological complexity decreases towards environment extremes. For example, ear size at silking is a major contributor to improved performance under drought (Cooper et al., 2014b; Messina et al., 2011; Messina et al., 2018). This is associated with silk elongation maintenance under water deficit (Hall et al., 1982; Fuad Hassan et al., 2008; Messina et al., 2019) and reduced ovule/kernel abortion (Edmeades et al., 1993; Messina et al., 2019). In contrast, in the absence of water deficit and ample nutrient availability, plant size and radiation use efficiency are major determinants of yield potential through radiation capture and transformation efficiency. At intermediate levels of productivity, which encompass most of the production environments in the US corn-belt, yield determination is dependent upon multiple traits and their interactions (gray area, Fig. 5), sometimes affecting yield in opposite ways, all occurring within a TPE, where environments oscillate from water deficit to mild water stress. Because it is possible to create a physiological profile for each hybrid in the selection set, it is also possible to apply the concept that each trait confers an advantage depending on the drought environment (Tardieu, 2012) to cope with uncertain weather. Breeders can select, and farmers can include in production systems hybrids that do not depend on the same mechanisms to determine yield in the target environments by accounting for the genetic correlations and the frequency of occurrence of drought environments. By doing so, it is feasible to deliver a set of hybrids that can increase the resilience of the production system by leveraging genetic and trait functional diversity. Biological knowledge, both genetic and physiological, can enable the creation of portfolios of products with improved yield stability as demonstrated by combining different crops in response to drought forecasts (Messina et al., 1999; Jones et al., 2000).

**Figure 5.**
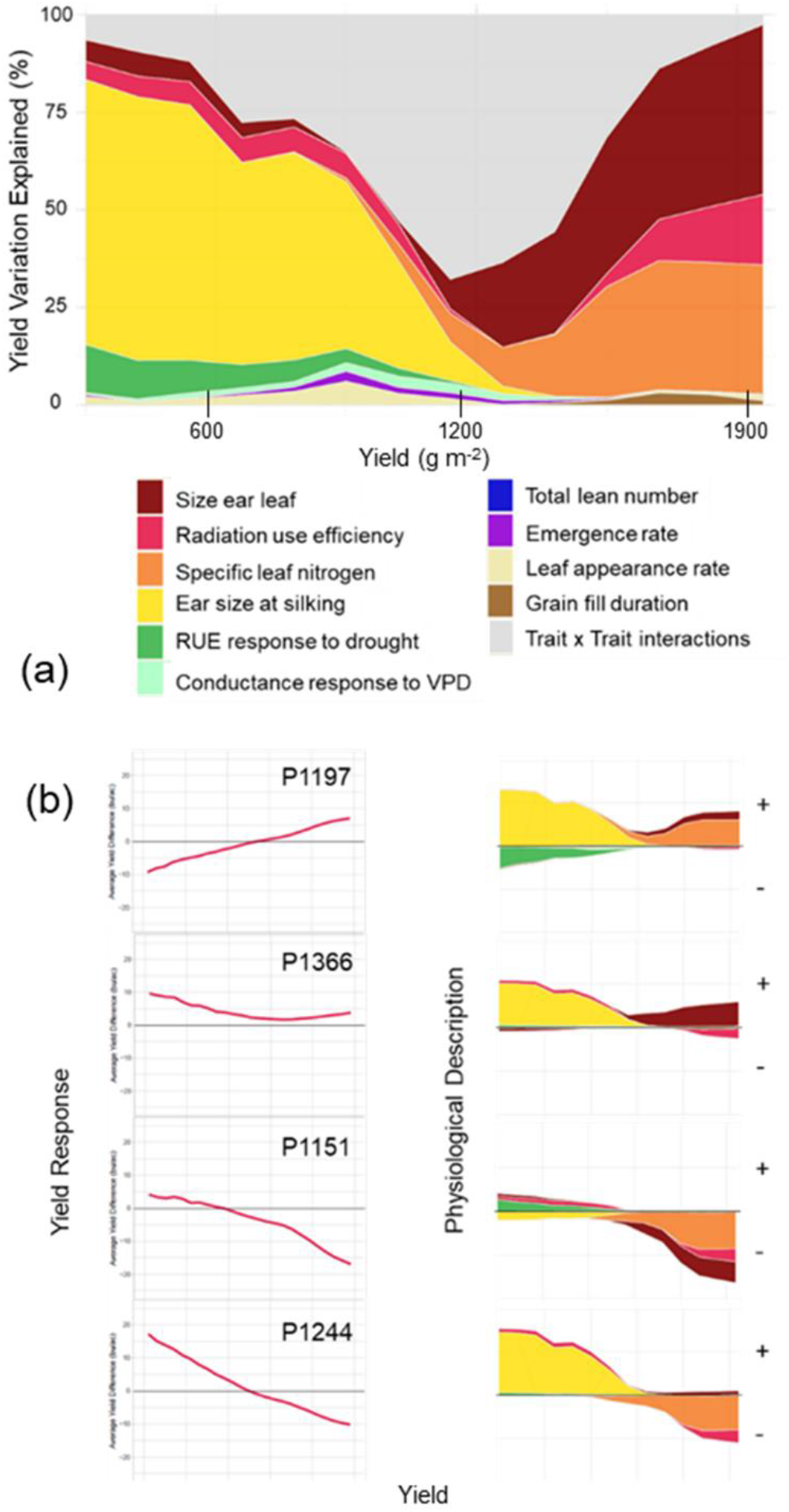
Biological determinants of yield across a range of environments represented as percent of total phenotypic variation (a), and examples for a set of contrasting hybrids along with a representation of yield variation with respect to average yield as a function of yield level (b).

Figure 5 shows that ear mass at silking is a major determinant of yield under water deficit and often discriminates DT from drought susceptible maize hybrids (Cooper et al., 2014b; Messina et al., 2018). To evaluate this prediction, data from a MET conducted at three locations in the central and western corn-belt in 2013 was analyzed for the relationship between yield and kernel set, and kernel set and ear size at silking. The experiment was conducted under rainfed conditions and managed according to the best management practices. Yield and kernel numbers were measured in three replicates at Johnston, IA, Garden City, KS, and Elgin, NE in three replicates, of four row plots spaced by 0.76 m and 5.4 m in length. Ear size at silking was measured at Johnston, IA on four plants and three replications. In this experiment, yield was highly associated with variation in kernel numbers across all three locations following the east-west precipitation gradient (Fig. 6). The correlations calculated between yield and ear size at silking by location across ten pre-commercial single cross hybrids were 0.6, 0.9, and 0.7 for Johnston, Elgin and Garden City, respectively. This result provides empirical evidence that conform well with the predictions from the biological model (Fig. 4, 5).

**Figure 6.**
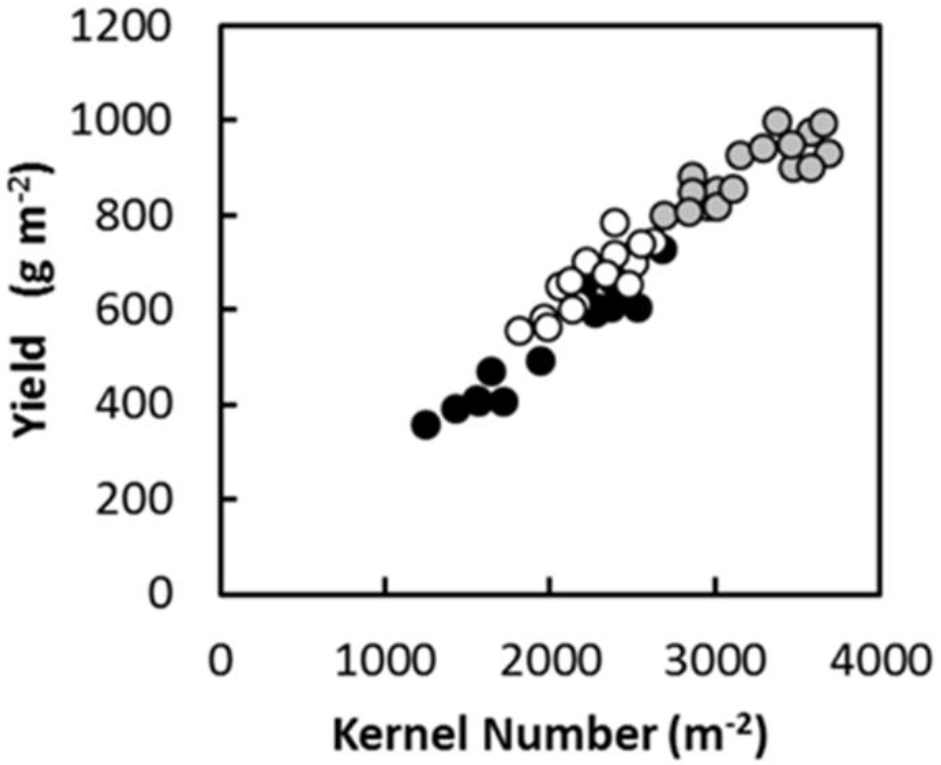
Yield under drought increased with increasing kernel set across an east-west precipitation gradient in the US Corn belt in 2013 and across experimental AQUAmax* hybrids. Johnston, IA 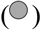, Elgin, NE (○) and Garden City, KS (●)

Models parameterized and evaluated as described above (Fig. 3, Fig. 4) can be utilized to inform selection and commercialization decisions (Fig. 1, 2). Figures 7 and 8 show an example of an ex-ante assessment of two commercial hybrids contrasting for their response to water deficit. Breeders and agronomists can assess the performance over thousands of virtual experiments across all of the US corn-belt and adjacent geographies for individual years or across a number of years of simulation. Figure 8 shows a clear domain of adaptation for the hybrids P1197 and P1244 with a transition point around 400 mm of water use. However, due to timing of rainfall and climate variability it is possible that for any field the performance could be reversed. Simulation results can quantify the probability for this to happen, thus creating opportunities to utilize not only information about the mean shift in yield but also the variability at a given location and potential management scenarios to inform decisions (Fig. 8). Producers’ attitudes towards risk are related to their degree of confidence when obtaining more precise information for deciding the best management practices with the goal of increasing the likelihood of improving yield and profits. Producers could be considered as slightly ‘risk-seeking’ with a mild degree of aversion (Bard and Barry, 2001). Frameworks that consider risk attitude can fully harness this information to manage climate risk through hybrid and crop diversification (Messina et al., 1999; Jones et al., 2000; Hammer et al. 2014, 2020).

**Figure 7.**
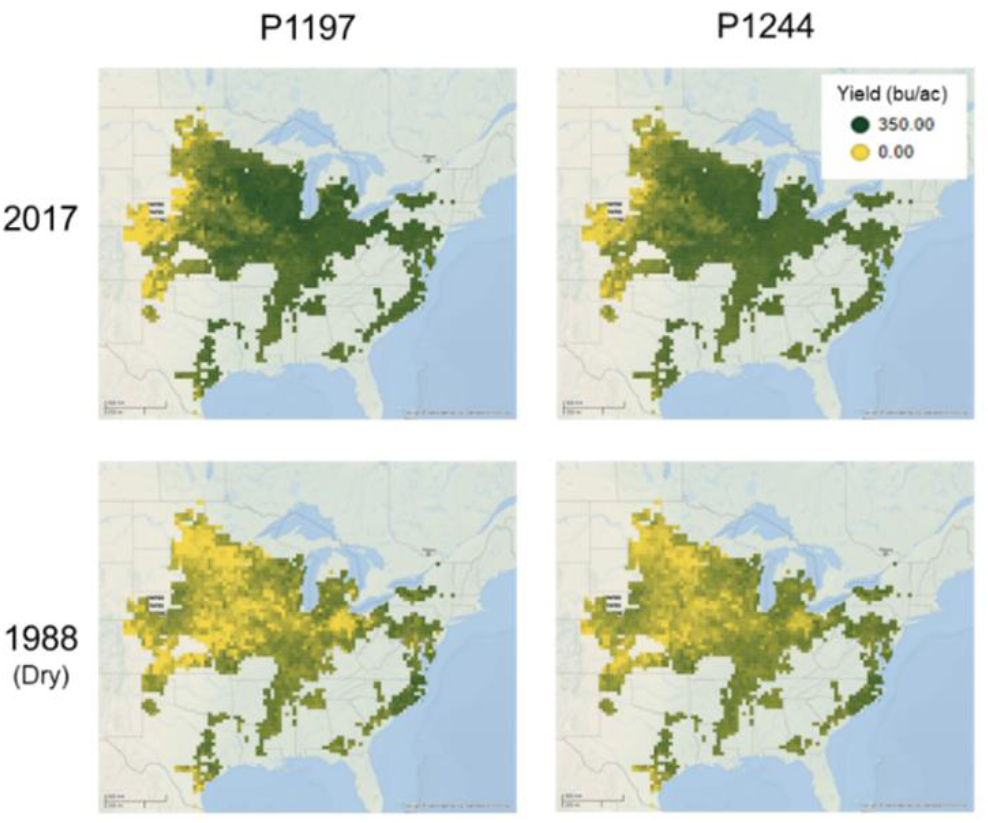
Simulated yields for two hybrids with contrasting behavior under water deficit for one dry year (1988), one wet year (2017) and 30 x 30 km grids characterized by a unique combination of soil, weather and agronomic management.

**Figure 8.**
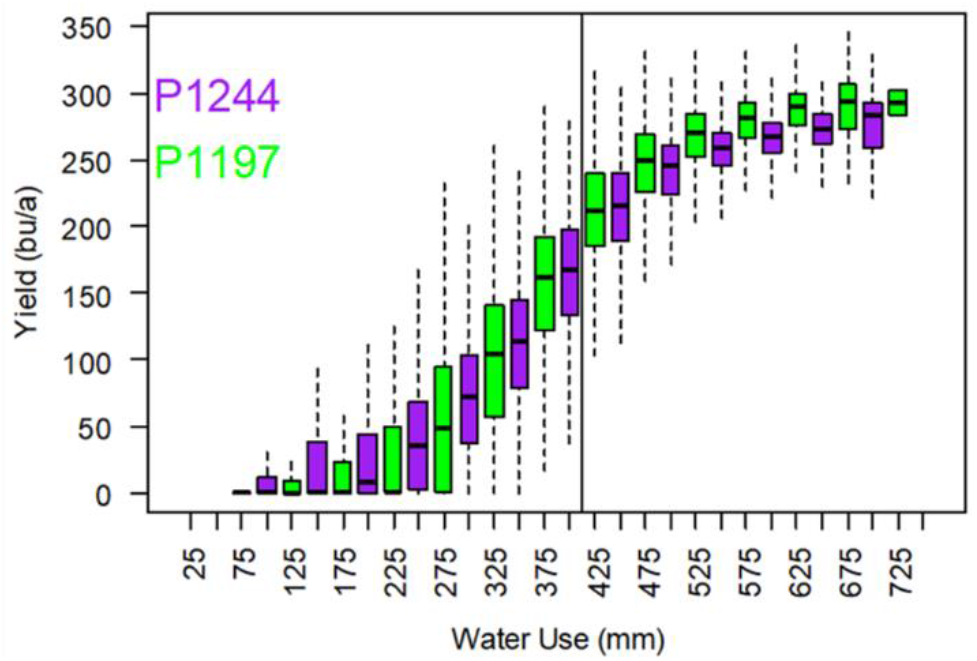
Simulated yield probability distributions along an environmental gradient defined by the water use for two hybrids contrasting in their response to water deficit.

### 2.3 On-Farm analyses and opportunities to improve dryland maize production using current and prospective genotype x management technologies

Predictive analytics for agronomic research and management of agricultural systems were available since the 1980s (Jones et al., 2017; Hammer et al., 2010). However, it was not until 2014 that we witnessed the application of prediction methods for on-farm analyses at scale for the use of N management in the central corn-belt. In contrast, only less than 1% of farmers use prediction and design methodologies to define strategies and manage irrigation in the United States (United States Department of Agriculture, 2019). Limited accessibility to the technology in user-friendly forms was implicated in explaining the low adoption. Instead, descriptive methodologies based on yield-ET empirical relations (Stone et al., 2006; Irmak et al., 2020) often inform irrigation planning and interpret crop productivity in rainfed systems (van Ittersum et al., 2013). Farmers incorporate hybrid DT scores to inform decisions, but the integration of the information is subjective. The empirical nature of yield-ET methods limits their applicability for prediction to an adjacent agronomic and genetic state space. These relationships are of great value to identify productivity gaps, but they need to be combined with prediction methods to enable design to close yield gaps through agricultural innovation (Cooper et al., 2020b; Messina et al., 2020b).

Only by harnessing biophysical and environment knowledge one can reimagine agricultural systems and conduct ex-ante evaluations to increase water productivity (Messina et al., 2018; Kruseman et al., 2020). While any trait can improve adaptation to drought stress in a particular context (Tardieu, 2012), the stochastic nature of the environmental processes that define repeatable and non-repeatable patterns of the TPE severely limits use of such awareness of dependency on context and our capacity to effectively sample the TPE. On-farm research programs could benefit from integrated approaches to prediction that account for properties of the TPE in the context of the germplasm available to the grower and their management practices. By combining genetic gain and yield gap methodologies, Cooper et al. (2020a) leveraged genetic and agronomic knowledge to transform descriptive yield-ET relations (Irmak et al., 2020) into a prediction framework (herein Digital Gap Analysis; DGA) to identify GxM technologies to close the production gap and increase water productivity across a range of water limited environments.

In contrast to yield-ET relations, the non-linear responses identified in DGA creates an opportunity to optimize water productivity for economically feasible yields. The DGA methodology provides quantitative and knowledge-based references for the grower to compare against alternative GxM technologies. The grower can optimize water productivity by 1) defining the ET level at which potential yield is attained, 2) identifying the domain within which ET and yield are linearly related, and 3) selecting GxM technologies to close the yield gaps. The discrepancy between linear and non-linear yield response to ET is explained by considering the linear response being one plausible realization out of many linear responses that could be observed within the GxExM state space defined by DGA, which implies that there are several possibilities to close yield gaps by leveraging GxM technologies.

Is the DGA framework applicable on-farm? To answer this question, an on-farm experiment with three producers in the western corn-belt was conducted using central pivot irrigation in 2019. Farms were in Webster County, NE, Chase County, NE and Thomas County, KS. The hybrid P1366 was planted under irrigation in the centers, and rainfed conditions in the corners of the fields with the center pivot with two replicates at each location. Yield was estimated using the farmers’ combines and ET was estimated from sixteen sensors deployed at each location using a modified surface renewal approach (McElrone et al., 2019). This method used semi-high frequency infrared radiometer surface temperature measurements to calculate sensible heat flux (H) to calculate latent heat flux (LE), and thereafter ET, as a residual to the energy budget R_n_= H + G + LE, where R_n_ is the net radiation and G is the soil heat flux.

Figure 9 shows the observed yields and ET pairs by environment and location. Quantile 99 and 80 are shown to quantify gaps. This result demonstrates the practical application of DGA on-farm. Using current technologies, it is feasible to implement systems to maximize water productivity. Overall, the results conform well with predictions from Cooper et al. (2020a). Minor differences in timing of irrigation (rainfed vs irrigated) for very similar ET levels led to a large productivity gap in Bluehill, NE in agreement with results from Cooper’s (2020) window experiment where reductions in irrigation were possible when water deficit was avoided at flowering time. These results suggest that in many of these rainfed environments supplemental deficit irrigation can have great impact on water productivity. Results are also consistent with the spread of simulated yields for a given level of ET (Cooper et al., 2020b; Fig. 8) and with the amplitude in observed yields shown by Cooper et al. (2020a) and Messina et al. (2019) in experiments conducted in managed-stress environments. The susceptibility of reproductive biology in maize (Daynard and Moldoon, 1983; Bolaños and Edmeades 1996) amplifies small differences in water deficit during the critical period for kernel set. The availability of image-based methods to estimate ET (McCabe and Wood, 2006; Jiang et al., 2020) can enable DGA for application at scale, and thus create avenues for improvement of water productivity. Estimates of ET can be obtained by using remote sensing-based models and satellite imagery data, with multiple methods and imagery data sources explored and tested during the last decade (Mu et al., 2009; Allen et al., 2011; Vyas et al., 2016; Yagci and Santanello, 2018). Considering the availability of phenotyping methods for ET, current opportunities to identify yield gaps at the farm scale (Fig. 9) and the ability to predict GxM (Fig. 3, Fig. 4), it is now possible to accelerate yield improvement and water productivity through selection.

**Figure 9.**
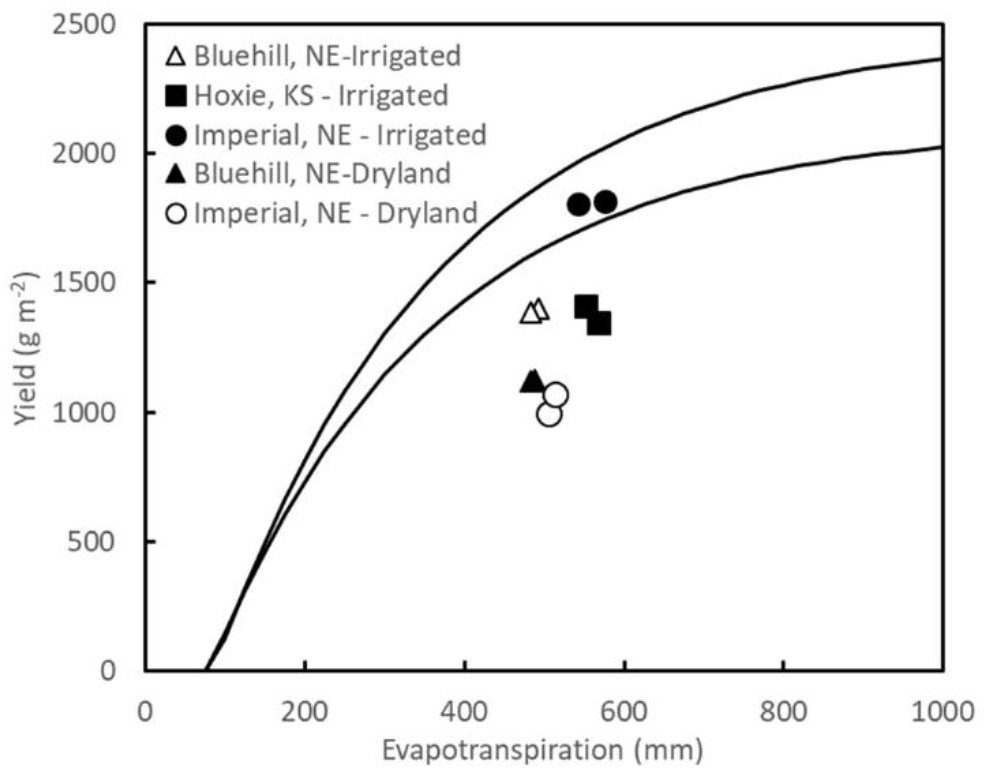
Theoretical maize yield response to evapotranspiration for quantiles 80 and 99 percentiles (lines) and yield observations for the hybrid P1366 at three locations in the western US corn belt for maize grown under rainfed and irrigated conditions, and under normal (closed symbols) and increased plant population by 1 pl m^−2^ (open symbols).

## 3 Design

### 3.1 Transitioning from evaluation to design of GxM technologies for cropping systems

Design is the user-centered process of imagining solutions to problems and the articulation of these in the form of blueprints that describe form and function of objects and systems that guide the subsequent process of creation. Design often must satisfy goals given a set of constraints. The use of optimization frameworks to find local or global solutions whenever possible was explored using tools to aid systems design (Peart and Curry,1998). As a user-focused activity, design starts by understanding the customer needs and with in-depth dialogues with agronomists. Often in product development, these product specifications take the form of a vector of thresholds for biotic and abiotic stress tolerance, standability and yield and yield stability metrics. With these blueprints, breeders and agronomists use phenotypic and genomic prediction to build a pipeline of biological products that minimizes the distance to targets (Cooper et al., 2014b). While the use of systems approaches to design solutions from field and regional scales dates back to the 1990s (Teng et al., 1997; Jones et al., 2000; Jones et al., 2017; Holzworth et al., 2014), the use of CGM in the design of crops is more recent (Hammer et al., 2014; Hammer et al., 2020; Cooper et al., 2020b). As part of a design toolkit, biophysical CGM and economic models were further integrated enabling ex-ante evaluation of designs and foresight analyses (Kruseman et al., 2020; Antle and Ray, 2020). Of importance for this review, are designs that seek to combine crops and genotypes in combinations with climate predictions to deal with climate risk and the devastating effects of drought (Messina, et al., 1999; Hammer et al., 2000; Jones et al., 2000). Current CGM with capabilities to utilize molecular markers for prediction, open opportunities to expand the decision set and augment the opportunities for farmers to cope with climatic risk through selection of DT genotypes among different crops.

During the last decade the assimilation of biological knowledge within models suitable for integration with genomic prediction was advanced (Hammer et al., 2010; Soufizadeh et al., 2018; Messina et al., 2019; Wu et al., 2019) and demonstrated for genetic improvement of drought tolerance in maize (Cooper et al., 2014b; Messina et al., 2018). This linked methodology created an unprecedented opportunity to harness genetics, agronomy and environmental science to enable design of GxM technologies to close crop improvement gaps identified at farm level (Fig., 9; Fig. 1). The debates about the required level of biological reality and approaches for integration of methodologies, however, have just begun (Hammer et al. 2019; Messina et al. 2020; Peng et al., 2020). Outcomes of this debate will enable translating genomic innovation (Bevan et al., 2017) into designs and breeding decisions. A range of approaches were proposed from the use of detailed mechanistic models (Hammer et al., 2019; Washburn et al., 2020) to the use of CGM to generate features to enhance prediction of statistical models (Boer et al., 2007; Rincent et al., 2019; van Eeuwijk et al., 2019; Washburn et al., 2020). Hammer et al. (2019) assessed the impact of changes of gene transformation on photo-biochemistry on yield across a range of environments using a biochemistry to field model based on APSIM (Wu et al., 2019). In contrast, Messina et al. (2018) demonstrated the use of simpler biological models embedded within a hierarchical Bayesian framework to improve genomic prediction in DT maize (Fig., 3,4). Despite the diversity of approaches, design blueprints could be now represented as vectors of markers, genes, physiological attributes, and agronomic management that could be contrasted with predicted scores between targets and genotypes that have never been tested. A more dynamic development of models is anticipated through iterative model building (Schrag, 1999; Messina et al., 2011; Hammer et al., 2019). The foundations of crop design have been established and provide the opportunity to build a new prediction-based paradigm for genetic improvement of crops.

#### Conclusions and perspectives

Molecular breeding approaches transformed breeding and dedicated efforts to improve DT in maize demonstrated sustained genetic gain at industrial scale and will continue providing the foundation to deliver DT maize. Here, our review demonstrates near-term opportunities to realize yield improvement that may include using technologies that harness both quantitative genetics and physiological frameworks for prediction at early stages of breeding, for placement of hybrids within regions, and design strategies given the DT hybrids and agronomic practices available to the grower. The feasibility to apply technologies to improve DT in maize from breeding to farm has the potential to accelerate crop improvement by designing and developing GxM technologies. DGA enables us to predict the outcome of combining haplotype genetic blocks that control physiological processes and agronomic practices even for genotypes that were created in a breeding program but never tested in the TPE. DGA in a way closes the cycle from breeding to farm and back to breeding.

Prediction methodologies were evolved and demonstrated to have the largest opportunities to deliver increased rates of crop improvement gain under water deficit conditions. Harnessing biological insights for end-to-end prediction is a promising path towards increasing yields and water productivity. However, there is a clear need for investments in plant science to advance our biological understanding of adaptation, germplasm diversity, algorithm development that improves statistical methodologies, and of most importance the development of a new engineering and design paradigm that harnesses complexity science and by doing so leverages noise and uncertainty to improve decisions and systems performance.

Crop growth model-whole genome prediction (CGM-WGP) methodology proved to be effective at modeling GxExM and has potential to improve decisions at all stages of product development and agriculture in drought prone environments. While the evaluation of CGM-WGP was specific to one combination of statistical and biophysical model, we argue that results could be generalized to the state that the combination of statistical learning and biological understanding can improve predictive skill. Model development and analytical approaches will be iterative as more information is gained through the process development pipeline and new data types are integrated. Closing the breeding-agronomy-production loop has potential to optimize both the effectiveness of the breeding program and farmers’ production.

## Acknowledgments

William S. Niebur, Geoff Graham, Jochen Scheel and John Arbuckle for their support over a decade of research. Andres Reyes and Andrea Salinas for their leadership at the Woodland and Viluco research stations. Chris Parry for his support to estimate ET on-farm.

## Abbreviations

AQ: AQUAmax®
ASI: anthesis-silking interval
CGM-WGP: crop growth model – whole genome prediction
DGA: Digital Gap Analyses
DT: drought tolerant maize
ET: evapotranspiration
MET: multi-environment trial
TPE: target population of environments
WGP: whole genome prediction

## References

Allen R, Irmak A, Trezza R, Hendrickx JMH, Bastiaanssen W, Kjaersgaard J. 2011. Satellite-based ET estimation in agriculture using SEBAL and METRIC. Hydrological Processes 25, 4011–4027.

Bevan MW, Uauy C, Wulff BBH, Zhou J, Krasileva K, Clark MD. 2017. Genomic innovation for crop improvement. Nature 543: 346–354.

Adee ED, Roozeboom K, Balboa GR, Schlegel A, Ciampitti IA. 2016. Drought-tolerant corn hybrids yield more in drought-stressed environments with no penalty in non-stressed environments. Frontiers in Plant Science, doi: 10.3389/fpls.2016.01534.

Antle JM, Ray S. 2020. Sustainable agricultural development: An economic perspective. Palgrave: McMillan.

Araus JL, Cairns JE. 2014. Field high-throughput phenotyping, the new frontier in crop breeding. Trends in Plant Science 19, 52–61.

Araus JL, Kefauver SC, Zaman-Allah M, Olsen MS, Cairns JE. 2018. Translating high-throughput phenotyping into genetic gain. Trends in Plant Science 23, 451–466.

Bard SK, Barry PJ. 2001. Assessing farmers’ attitudes toward risk using the “Closing-in” method. Journal of Agricultural and Resource Economics 26, 248–260.

Barker T, Campos H, Cooper M, Dolan D, Edmeades G, Habben J, Schussler J, Wright D, Zinselmeier C. 2005. Improving drought tolerance in maize. Plant Breeding Reviews 25, 173–253.

Boer, MP, Wright D, Feng L, Podlich DW, Luo L, Cooper M and van Eeuwijk FA. 2007. A Mixed-Model Quantitative Trait Loci (QTL) Analysis for Multiple-Environment Trial Data Using Environmental Covariables for QTL-by-Environment Interactions, With an Example in Maize. Genetics 77, 1801–1813.

Bogard M, Biddulph B, Zheng B, Hayden M, Kuchel H, Mullan D, Allard V, Le Gouis J, Chapman SC. 2020. Linking genetic maps and simulation to optimize breeding for wheat flowering time in current and future climates. Crop Science 60, 678–699.

Bolaños J, Edmeades GO. 1996. The importance of the anthesis-silking interval in breeding for drought tolerance in tropical maize. Field Crops Research 48, 65–80.

Boyer JS, Byrne P, Cassman KG, et al. 2013. The US drought of 2012 in perspective: a call to action. Global Food Security 2, 139–143.

Bruce WB, Edmeades GO, Barker TC. 2002. Molecular and physiological approaches to maize improvement for drought tolerance. Journal of Experimental Botany 53, 13–25.

Bustos-Korts D, Boer MP, Malosetti M, Chapman S, Chenu K, Zheng B, van Eeuwijk FA. 2019a. Combining crop growth modeling and statistical genetic modeling to evaluate phenotyping strategies. Frontiers in Plant Science doi: 10.3389/fpls.2019.01491.

Bustos-Korts D, Malosetti M, Chenu K, Chapman S, Boer MP, Zheng B, van Eeuwijk FA. 2019b. From QTLs to adaptation landscapes: Using genotype-to-phenotype models to characterize GxE over time. Frontiers in Plant Science doi: 10.3389/fpls.2019.01540.

Campos H, Cooper M, Habben JE, Edmeades GO, Schussler JR. 2004. Improving drought tolerance in maize: a view from industry. Field Crops Research 90, 19–34.

Casadebaig P, Bangyou Z, Chapman S, Huth N, Faivre R, Chenu K. 2016. Assessment of the potential impacts of wheat plant traits across environments by combining crop modeling and global sensitivity analysis. PLoS One 11, e0146385.

Castiglioni P, Warner D, Bensen RJ, et al. 2008. Bacterial RNA chaperones confer abiotic stress tolerance in plants and improved grain yield in maize under water-limited conditions. Plant Physiology 147, 446–455.

Chapman S, Cooper M, Podlich D, Hammer G. 2003. Evaluating plant breeding strategies by simulating gene action and dryland environment effects. Agronomy Journal 95, 99–113.

Choudhary S, Sinclair TR, Messina CD, Cooper M. 2013. Hydraulic conductance in maize hybrids differing in breakpoint of transpiration response to increasing vapor pressure deficit. Crop Science 54, 1147–1152.

Cooper M, Gho C, Leafgren R, Tang T, Messina C. 2014a. Breeding drought-tolerant maize hybrids for the US corn-belt: discovery to product. Journal of Experimental Botany 65, 6191–6204.

Cooper M, Messina CD, Podlich D, Totir LR, Baumgarten A, Hausmann NJ, Wright D, Graham G. 2014b. Predicting the future of plant breeding: complementing empirical evaluation with genetic prediction. Crop and Pasture Science 65, 311–336.

Cooper M, Powell O, Voss-Fels KP, Messina CD, Gho C, Podlich DW, Technow F, Chapman SC, Beveridge CA, Ortiz-Barientos D, Hammer GL. 2020c. Modelling selection response in plant breeding programs using crop models as mechanistic gene-to-phenotype (CGM-G2P) multi-trait link functions. doi.org/10.1101/2020.10.13.338301.

Cooper M, Tang T, Gho C, Hart T, Hammer G, Messina C. 2020a. Integrating Genetic Gain and Gap Analysis to predict improvements in crop productivity. Crop Science 60, 582–604.

Cooper M, Voss-Fels KP, Messina CD, Tang T, Hammer GL. 2020b. Tackling GxExM interactions to close on-farm yield-gaps: Creating novel pathways for crop improvement by predicting contributions of genetics and management to crop productivity. Theoretical Applied Genetics (in review)

Crossa J. Pérez-Rodríguez P, Cuevas J et al. 2017. Genomic selection in plant breeding: methods, models, and perspectives. Trends in Plant Science 22, 961–975.

Daynard TB, Moldoon JF. 1983. Plant-to-plant variability of maize plants grown at different densities. Canadian Journal of Plant Science 63, 45–59.

Debruin J, Messina CD, Munaro E, Thompson K, Conlon-Beckner C, Fallis L, Sevenich DM, Gupta R, Dhugga KS. 2013. N distribution in maize plant as a marker for grain yield and limits on its remobilization after flowering. Plant Breeding 132, 500–505.

Edmeades GO, Hernandez JBM, Bello S. 1993. Causes for silk delay in a lowland tropical maize population. Crop Science 33, 1029–1035.

Fuad Hassan A, Tardieu F, Turc O. 2008. Drought induced changes in anthesis-silking interval are related to silk expansion: a spatio-temporal growth analysis in maize plants subjected to soil water deficit. Plant Cell and Environment 31, 1349–136.

Fraser AS, Burnell DG. 1970. Computer models in Genetics. McGraw-Hill, San Francisco, CA.

Gaffney J, Schussler J, Löffler C, Cai W, Paszkiewicz S, Messina C, Groetke J, Keaschall J, Cooper M. 2015. Industry-scale evaluation of maize hybrids selected for increased yield in drought-stress conditions of the US corn belt. Crop Science 55:1608–1618.

Gianola D, de los Campos G, Hill WG, Manfredi E, Fernando R. 2009. Additive genetic variability and the Bayesian alphabet. Genetics 183, 347–363.

Guo M, Rupe MA, Wei J, et al. 2014. Maize ARGOS1 (ZAR1) transgenic alleles increase hybrid maize yield. Journal of Experimental Botany 65, 249–260.

Habben JE, Bao X, Bate NJ, DeBruin JL, Dolan D, Hasegawa D, Helentjaris TG, Lafitte RH, Lovan N, Mo H, Reimann K, Schussler JR. 2014. Transgenic alteration of ethylene biosynthesis increases grain yield in maize under field drought-stress conditions. Plant Biotechnology Journal 12, 685–693.

Hall AJ, Vilella F, Trapani N, Chimenti C. 1982. The effects of water stress and genotype on the dynamics of pollen-shedding and silking in maize. Field Crops Research 5, 349–363.

Hammer GL, Dong Z, McLean G, Doherty A, Messina C, Schusler J, Zinselmeier C, Paszkiewicz S, Cooper M. 2009. Can changes in canopy and/or root system architecture explain historical maize yield trends in the US Corn Belt? Crop Science 49, 299–312.

Hammer GL, McLean G, Chapman S, Zheng B, Doherty A, Harrison MT, van Oosterom E, Jordan D. 2014. Crop design for specific adaptation in variable dryland production environments. Crop and Pasture Science 65, 614–626.

Hammer GL, McLean G, van Oosterom E, Chapman S, Zheng B, Wu A, Doherty A, Jordan D. 2020. Designing crops for adaptation to the drought and high-temperature risks anticipated in future climates. Crop Science 60, 605–621.

Hammer G, Messina C, Wu A, Cooper M. 2019. Biological reality and parsimony in crop models – why we need both in crop improvement! in silico Plants 1:diz010

Hammer GL, Nicholls N, Mitchell C. 2000. Applications of Seasonal Climate Forecasting in Agricultural and Natural Ecosystems. Springer.

Hammer GL, van Oosterom E, McLean G, Chapman SC, Broad I, Harlan P, Muchow RC. 2010. Adapting APSIM to model the physiology and genetics of complex adaptive traits in field crops. Journal of Experimental Botany 61, 2185–2202.

Hao B, Xue Q, Marek TH, et al. 2015a. Water use and grain yield in drought-tolerant corn in the Texas high plains. Agronomy Journal 107, 1922–1930.

Hao B, Xue Q, Marek TH, et al. 2019. Grain yield, evapotranspiration, and water-use efficiency of maize hybrids differing in drought tolerance. Irrigation Science 37, 25–34.

Hao B, Xue Q, Marek TH, Jessup KE, Hou X, Xu W, Bynum ED, Bean BW. 2015b. Soil water extraction, water use, and grain yield by drought-tolerant maize on the Texas High Plains. Agricultural Water Management 155, 11–21.

Heslot N, Akdemir D, Sorrells ME, Jannink JL. 2014. Integrating environmental covariates and crop modeling into the genomic selection framework to predict genotype by environment interactions. Theoretical Applied Genetics 127, 463–480.

Holzworth, DP, Huth NI, Devoil PG, et al. 2014. APSIM-Evolution towards a new generation of agricultural systems simulation. Environmental Modelling and Software 62, 327–350.

Hochholdinger F, Yu P, Marcon C. 2018. Genetic control of root system development in maize. Trends Plant Science 23, 79–88.

Jarquin D, Crossa J, Lacaze X, et al. 2014. A reaction norm model for genomic selection using high-dimensional genomic and environmental data. Theoretical Applied Genetics 127, 595–607.

Jarquín D, Lemes da Silva C, Gaynor RC, Poland J, Fritz A, Howard R, Battenfield S, Crossa J. 2017. Increasing genomic-enabled prediction accuracy by modeling genotype × environment interactions in Kansas wheat. The Plant Genome, 10: 1–15

Jiang C, Guan K, Pan M, Ryu Y, Peng B, Wang S. 2020. BESS-STAIR: a framework to estimate daily, 30 m, and all-weather crop evapotranspiration using multi-source satellite data for the US Corn Belt. Hydrology and Earth System Sciences 24, 1251–1273.

Jones JW, Antle JM, Basso B, et al. 2017. Brief history of agricultural systems modeling. Agricultural Systems 155, 240–254.

Jones JW, Hansen JW, Royce FS, Messina, CD. 2000. Potential benefits of climate forecasting to agriculture. Agriculture, Ecosystems and Environment 82, 169–184.

Kruseman G, Bairagi S, Komarek, AM, A Molero Milan, Nedumaran S, Petsakos A, Prager S, Yigezu YA. 2020. CGIAR modeling approaches for resource-constrained scenarios: II. Models for analyzing socioeconomic factors to improve policy recommendations. Crop Science 60, 568–581.

Lacube S, Fournier C, Palaffre C, Millet EJ, Tardieu F, Parent B. 2017. Distinct controls of leaf widening and elongation by light and evaporative demand in maize. Plant, Cell and Environment 40, 2017–2028.

Li X, Guo T, Yu J. 2018. Genomic and environmental determinants and their interplay underlying phenotypic plasticity. Proceedings National Academy of Science 115, 6679–6684.

Lindsey AJ, Thomison PR. 2016. Drought-tolerant corn hybrid and relative maturity yield response to plant population and planting date. Agronomy Journal 108, 229–242.

Lynch M, Walsh B. 1998. Genetics and Analysis of Quantitative Traits. Sinauer Associates, Inc., Sunderland, MA.

Lowenberg-DeBoer J, Erickson B. 2019. Setting the Record Straight on Precision Agriculture Adoption. Agronomy Journal 111, 1552–1569.

McCabe MF, Wood EF. 2006. Scale influences on the remote estimation of evapotranspiration using multiple satellite sensors. Remote Sensing of Environment 105, 271–285.

McCormick RF, Truong SK, Rotundo J, Gaspar AP, Kyle D, van Eeuwijk F, Messina CD. 2020. Intercontinental prediction of soybean phenology via hybrid ensemble of knowledge-based and data-driven models doi: https://doi.org/10.1101/2020.09.22.30650

McElrone AJ, Bambach-Ortiz NE, Parry CK. 2019. An alternative method to estimate atmosphere-canopy fluxes from semi-high frequency canopy infrared temperature. AGUFM. Dec;2019:B31N–2399.

McFadden J, Smith D, Wechsler S, Wallander S. 2019. Development, adoption, and management of drought-tolerant corn in the United States. EIB-204, U.S. Department of Agriculture, Economic Research Service.

Millet E, Kruijer W, Coupel-Ledru A, et al. 2019. Genomic prediction of maize yield across European environmental conditions. Nature Genetics 51, 952–956.

Messina C, Cooper M, McDonald D, Poffenbarger H, Clark R, Salinas A, Fang Y, Gho C, Tang T, Graham G. 2020a. Reproductive resilience but not root architecture underpin yield improvement in maize. doi.org/10.1101/2020.09.30.320937

Messina C, Cooper M, Reynolds M, Hammer G. 2020b. Crop science: A foundation for advancing predictive agriculture. Crop Science 60, 544–546.

Messina CD, Hammer GL, McLean G, Cooper M, van Oosterom EJ, Tardieu F, Chapman SC, Doherty A, Gho C. 2019. On the dynamic determinants of reproductive failure under drought in maize. in silico Plants, doi: 10.1093/insilicoplants/diz003

Messina CD, Hansen JW, Hall AJ. 1999. Land allocation conditioned on El Niño-Southern Oscillation phases in the pampas of Argentina. Agricultural Systems 60, 197–212.

Messina CD, Jones JW, Boote KJ, Vallejos CE. 2006 A Gene-based model to simulate soybean development and yield responses to environment. Crop Science 46, 456–466.

Messina CD, Podlich D, Dong Z, Samples M, Cooper M. 2011. Yield–trait performance landscapes: from theory to application in breeding maize for drought tolerance. Journal of Experimental Botany 62, 855–868.

Messina CD, Sinclair TR, Hammer GL, Curan D, Thompson J, Oler Z, Gho C, Cooper M. 2015. Limited-Transpiration trait may increase maize drought tolerance in the US Corn Belt. Agronomy Journal 107:1978–1986.

Meuwissen THE, Hayes BJ, Goddard ME. 2001. Prediction of total genetic value using genome-wide dense marker maps. Genetics 157, 1819–1829.

Mounce RB, O’Shaughnessy SA, Blaser BC, Colaizzi PD, Evett SR. 2016. Crop response of drought-tolerant and conventional maize hybrids in a semiarid environment. Irrigation Science 34, 231–244.

Mu Q, Jones LA, Kimball JS, McDonald KC, Running SW. 2009. Satellite assessment of land surface evapotranspiration for the pan-Arctic domain. Water Resources Research 45, W09420. doi:10.1029/2008WR007189.

Pathak TB, Fraisse CW, Jones JW, Messina CD, Hoogenboom G. 2007. Use of global sensitivity analysis for CROPGRO cotton model development. Transactions of the ASABE 50, 2295–2302.

Peart RM, Curry RB. 1998. Agricultural systems modeling and simulation. Marcel Dekker.

Peng BK, Guan J, Tang EA, et al. 2020. Advancing multiscale crop modeling for agricultural climate change adaptation assessment. Nature Plants 6, 338–348.

Podlich DW, Cooper M. 1998. QU-GENE: A simulation platform for quantitative analysis of genetic models. Bioinformatics 14, 632–653.

Podlich DW, Cooper M, Basford KE. 1999. Computer simulation of a selection strategy to accommodate genotype-environment interactions in a wheat recurrent selection programme. Plant Breeding 118, 17–28.

Ramirez-Villegas J, Molero Milan A, Alexandrov N, et al. 2020. CGIAR modeling approaches for resource-constrained scenarios: I. Accelerating crop breeding for a changing climate. Crop Science 60, 547–567.

Reyes A, Messina CD, Hammer GL, Liu L, van Oosterom E, Lafitte R, Cooper M. 2015. Soil water capture trends over 50 years of single-cross maize (Zea mays L.) breeding in the US corn-belt. Journal of Experimental Botany 66, 7339–7346.

Reynolds M, Chapman S, Crespo-Herrera L, Molero G, Mondal S, Pequeno DNL, Pinto F, Pinera-Chavez FJ, Poland J, Rivera-Amado C, Saint Pierre C, Sukumaran S. 2020. Breeder friendly phenotyping. Plant Science 295: 110396. https://doi.org/10.1016/j.plantsci.2019.110396

Rincent R, Malosetti M, Ababaei B, Touzy G, Mini A, Bogard M, Martre P, Le Gouis J, van Eeuwijk F. 2019. Using crop growth model stress covariates and AMMI decomposition to better predict genotype-by-environment interactions. Theoretical Applied Genetics 132, 3399–3411.

Schrag M. 1999. Serious play: how the world’s best companies simulate to innovate. Harvard Business School Press.

Shekoofa A, Sinclair TR, Messina CD, Cooper M. 2015. Variation among maize hybrids in response to high vapor pressure deficit at high temperatures. Crop Science 55, 392–396.

Shi J, Gao H, Wang H, Lafitte HR, Archibald RL, Yang M, Hakimi SM, Mo H, Habben JE. 2017. ARGOS8 variants generated by CRISPR-Cas9 improve maize grain yield under field drought stress conditions. Plant Biotechnology Journal 15, 207–216.

Shi J, Habben JE, Archibald RL, Drummond BJ, Chamberlin MA, Williams RW, Lafitte HR, Weers BP. 2015. Overexpression of ARGOS genes modifies plant sensitivity to ethylene, leading to improved drought tolerance in both arabidopsis and maize. Plant Physiology 169, 266–282.

Soufizadeh S, Munaro E, McLean G, Massignam A, van Oosterom EJ, Chapman SC, Messina C, Cooper M, Hammer GL. 2018. Modelling the nitrogen dynamics of maize crops – Enhancing the APSIM maize model. European Journal of Agronomy 100, 118–131.

Ruta N, Liedgens M, Fracheboud Y, Stamp P, Hund A. 2010. QTLs for the elongation of axile and lateral roots of maize in response to low water potential. Theoretical and Applied Genetics 120, 621–631.

Stone LR, Schlegel AJ, Khan AH, Klocke NL, Aiken RM. 2006. Water supply: yield relationships developed for study of water management. Journal of Natural Resources and Life Science Education 35, 161–173.

Tanner CB, Sinclair TR. 1983. Efficient water use in crop production: Research or research? In: Taylor HM, Jordan WR, editors, Limitations to efficient water use in crop production. ASA, CSSA, and SSSA, Madison, WI. p. 1–27.

Tardieu F. 2012. Any trait or trait-related allele can confer drought tolerance: just design the right drought scenario. Journal of Experimental Botany 63, 25–31.

Tardieu F, Simonneau T, Parent B. 2017. Modelling the coordination of the controls of stomatal aperture, transpiration, leaf growth, and abscisic acid: update and extension of the Tardieu–Davies model. Journal of Experimental Botany 66, 2227–2237.

Teng PS, Kropff MJ, ten Berge HFM, Dent JB, Lansigan FP, van Laar HH. 1997. Application of systems approaches at the farm and regional levels. Vol 1. Kluwer Academic Publishers.

Tuberosa R, Sanguineti MC, Landi PL, Giuliani MM, Salvi S, Conti S. 2002. Identification of QTLs for root characteristics in maize grown in hydroponics and analysis of their overlap with QTLs for grain yield in the field at two water regimes. Plant Molecular Biology 48: 697–712.

Turc O, Bouteille M, Fuad-Hassan A, Welcker C, Tardieu F. 2016. The growth of vegetative and reproductive structures (leaves and silks) respond similarly to hydraulic cues in maize. New Phytologist 212, 377–388.

United States Department of Agriculture. 2019. 2018 Irrigation and water management survey. Volume 3. Special Studies. Part 1. AC-17-SS-1

van Ittersum MK, Cassman KG, Grassini P, Wolf J, Tittonelli P, Hochman Z. 2013. Yield gap analyses with local to global relevance—A Review. Field Crops Research 143, 4–17.

van Eeuwijk FA, Bustos-Korts D, Millet EJ, et al. 2019. Modelling strategies for assessing and increasing the effectiveness of new phenotyping techniques in plant breeding. Plant Science 282, 23–39.

van Eeuwijk FA, Boer ML, Totir R, et al. 2010. Mixed model approaches for the identification of QTLs within a maize hybrid breeding program. Theoretical and Applied Genetics 120, 429–440.

van Oosterom EJ, Yang Z, Zhang F, Deifel KS, Cooper M, Messina CD, Hammer GL. 2016. Hybrid variation for root system efficiency in maize: potential links to drought adaptation. Functional Plant Biology 43, 502–511.

Voss-Fels KP, Cooper M, Hayes BJ. 2019. Accelerating crop genetic gains with genomic selection. Theoretical and Applied Genetics 132, 669–686.

Vyas SS, Nigam R, Bhattacharya BK, Kumar P. 2016. Development of real-time reference evapotranspiration at the regional scale using satellite-based observations. International Journal of Remote Sensing 37, 6108–6126. doi:10.1080/01431161.2016.1253895.

Wallach D, Makowski D, Jones JW, Brun F. 2019. Working with dynamic crop models. 3rd edition. Academic Press.

Walsh B, Lynch M. 2018. Evolution and Selection of Quantitative Traits. Sinauer Associates, Oxford University Press, Oxford.

Washburn JD, Burch MB, Franco JAV. 2020. Predictive breeding for maize: Making use of molecular phenotypes, machine learning, and physiological crop models. Crop Science 60, 622–638.

Wu A, Hammer GL, Doherty A, von Caemmerer, Farquhar GD. 2019. Quantifying impacts of enhancing photosynthesis on crop yield. Nature Plants 5, 380–388.

Yagci AL, Santanello JA. 2018. Estimating evapotranspiration from satellite using easily obtainable variables: A case study over the southern great plains, U.S.A. IEEE Journal of Selected Topics in Applied Earth Observations and Remote Sensing 11, 12–23. doi: 10.1109/JSTARS.2017.2753723.

Yin X, Struik PC, Kropff MJ. 2004. Role of crop physiology in predicting gene-to-phenotype relationships. Trends Plant Science 9, 426–432.

Zhao J, Xue Q, Jessup KE, Hao B, Hou X, Marek TH, Xu W, Evett SR, O’Shaughnessy SA, Brauer DK. 2018. Yield and water use of drought-tolerant maize hybrids in a semiarid environment. Field Crops Research 216, 1–9.

